# Age-Related Modulations of Alpha and Gamma Brain Activities Underlying Anticipation and Distraction

**DOI:** 10.1101/496117

**Authors:** Hesham A. ElShafei, Lesly Fornoni, Rémy Masson, Olivier Bertrand, Aurélie Bidet-Caulet

## Abstract

Attention operates through top-down (TD) and bottom-up (BU) mechanisms. Recently, it has been shown that slow (alpha) frequencies index facilitatory and suppressive mechanisms of TD attention and faster (gamma) frequencies signal BU attentional capture. Ageing is characterized by increased behavioral distractibility, resulting from either a reduced efficiency of TD attention or an enhanced triggering of BU attention. However, only few studies have investigated the impact of ageing upon the oscillatory activities involved in TD and BU attention. MEG data were collected from 14 elderly and 14 matched young healthy human participants while performing the Competitive Attention Task. Elderly participants displayed (1) exacerbated behavioral distractibility, (2) altered TD suppressive mechanisms, indexed by a reduced alpha synchronization in task-irrelevant regions, (3) less prominent alpha peak-frequency differences between cortical regions, (4) a similar BU system activation indexed by gamma activity, and (5) a reduced activation of lateral prefrontal inhibitory control regions. These results show that the ageing-related increased distractibility is of TD origin.

## 1 Introduction

Imagine being in a game of bingo, waiting for the next number to be called out, and suddenly your phone starts ringing. In such situations, our brains rely on a balance between top-down (TD) anticipatory attentional mechanisms to better process upcoming (relevant) stimuli and bottom-up (BU) attentional capture that allows the processing and evaluation of distracting (irrelevant) stimuli (1). In this context, on one hand, TD attention promotes the processing of relevant stimuli through facilitatory and suppressive mechanisms, resulting in enhanced processing of relevant information and reduced brain responses to unattended inputs, respectively (review in 2–6). On the other hand, BU attention refers to the attentional capture by task-irrelevant unexpected salient stimuli (5, 7). This apparent dichotomy does not exclude the deployment of other attentional systems during different tasks or situations (8).

A good balance between the aforementioned TD and BU mechanisms is thus crucial: enhanced distractibility can result from either a reduced efficiency of TD mechanisms, or an enhanced triggering of BU attentional capture. TD and BU processes are supported by partially segregated networks: (i) a dorsal fronto-parietal network, including the intraparietal sulcus and the frontal eye fields, and (ii) a ventral fronto-parietal network, including the temporo-parietal junction and the ventral frontal cortex, respectively. The two networks mainly overlap in the lateral prefrontal cortex (lPFC) (1,9,10). Indeed, it has been demonstrated that the lPFC could be involved in the top–down control of attention by means of biasing sensory processing in favor of information that is behaviorally relevant (11–13), and by mediating the inhibition of distracting stimulus processing (e.g. 14).

Moreover, it has been suggested that neural communication (signal transmission) within these networks could propagate via distinct frequency bands with slower frequencies, namely alpha, supporting interactions along the TD network and faster frequencies, namely gamma, supporting communication within the BU network (15–17). Importantly this dichotomy between slow and fast oscillatory activities is not specific to attentional processes, as it stems from a line of studies demonstrating that slow and fast oscillations would specifically support feedback and feedforward neural communication, respectively (15,17–21).

Alpha oscillations (8–14 Hz) have been proposed to play a crucial role in TD anticipatory attention (review in 22,23, but see 24). More precisely, they play an active inhibitory role (25, 26): reduced and enhanced alpha power reflect increased and decreased cortical excitability in task relevant and irrelevant areas, respectively (e.g. 27–30). Therefore, alpha rhythm is a suitable candidate for supporting facilitatory and suppressive mechanisms of anticipatory attention (6). Importantly, in a recent study, we could show that that facilitatory and suppressive attention mechanisms are supported by different alpha sub-bands (6). Gamma oscillations (>30 Hz) have also been associated with attention (review in 31) with evidence suggesting that BU feedforward signaling propagates pre-dominantly via these oscillations in primate sensory areas (32, 33). Moreover, gamma activity has been found in frontal regions of the ventral network in response to novel sounds (34–36).

Ageing is characterized by attentional difficulties, in particular, a reduced capability to inhibit irrelevant information (37, 38). This exacerbated distractibility has been attributed to a degradation of inhibitory mechanisms (inhibitory deficit hypothesis, 39) and/or a deterioration in the frontal lobe functioning (frontal aging hypothesis, 40). With ageing, TD attentional facilitatory processes, as indexed by alpha desynchronization, have been found to be either reduced (41–43), preserved (44) or even enhanced (45). However, TD suppressive processes indexed by alpha synchronization seem to deteriorate (46). Moreover, to our knowledge, no study has investigated the impact of ageing on gamma oscillatory activity supporting BU attention.

Thus, the aim of the present study was to characterize the brain origins of the exacerbated distractibility in the elderly by investigating the impact of ageing on oscillatory activities supporting the balance between TD and BU attention. For this purpose, we recorded MEG activity from young and elderly participants while performing the Competitive Attention Task (47), a novel paradigm that permits the assessment of BU and TD mechanisms of auditory attention and the interaction between them. To assess voluntary attention orienting, this task includes informative and uninformative visual cues respectively indicating - or not - the spatial location of a forthcoming auditory target to detect. To measure distraction, the task comprises trials with a task-irrelevant distracting sound preceding the target according to several delays. This change in distractor timing onset allows to dissociate the effects of distraction and phasic arousal triggered by a distracting sound (47–54).

We hypothesized that ageing would be characterized by an (1) exacerbated behavioral distractibility, (2)reduced TD filtering of irrelevant information, indexed by alterations in the alpha band, and (3) altered gamma responses to distracting sounds.

## 2 Material & Methods

### 2.1 Participants

Participants were 14 young (mean age = 25 ± 0.67 Standard Error of Mean (SEM); range: 20-29 years; 5 females) and 14 elderly (mean age = 67 ± 1.08 SEM; range: 61-75 years; 5 females) adults. The two groups were matched for sex, handedness, scholar and musical education (see Table 1). As expected, the 2 groups significantly differ in age (unpaired t-test p < 0.001) but did not significantly differ in scholar (unpaired t-test p = 0.67) and musical (unpaired t-test p = 0.32) education. All participants were healthy, right-handed, free from any neurological or psychiatric disorders and reported normal hearing, and normal or corrected-to-normal vision. Data regarding scholarship and musical education were collected by asking the participants the latest educational degree obtained and the number of years of formal musical training, respectively. In addition, all participants performed the Mini-Mental State Examination (MMSE, 55) in order to compare cognitive abilities between young and elderly participants and no significant difference was found (unpaired t-test p = 0.52). The study was approved by the local ethical committee (CPP Sud-Est III, authorization number 2014-050B), and subjects gave written informed consent, according to the Declaration of Helsinki, and they were paid for their participation. Please note that data from all young participants are included in the analysis presented in a previous study of gamma activity in young adults (36).

**Table 1.** Group demographics. SEM, standard error of the mean; F, Female; M, Male; R, right-handed.

|  | **Elderly group** | **Young Group** | **Unpaired t-test**<br>**p value** |
| --- | --- | --- | --- |
| **Age (years ± SEM)** | 67 ± 1.08 | 25 ± 0.67 | < 0.001 |
| **Gender** | 5F, 8M | 5F, 8M | NA |
| **Handedness** | 14R | 14R | NA |
| **Scholar Education (years ± SEM)** | 15 ± 0.71 | 15 ± 0.57 | 0.67 |
| **Musical Education (years ± SEM)** | 1 ± 0.45 | 2 ± 0.68 | 0.32 |
| **MMSE (± SEM)** | 29.21 ± 0.29 | 28.85 ± 0.46 | 0.52 |

### 2.2 Stimuli and tasks

#### 2.2.1 Competitive Attention Task (CAT)

In all trials, a central visual cue (200 ms duration) indicated the location of a target sound (100 ms duration) that followed with a fixed delay of 1000 ms (see Figure 1). The cue was a green arrow, presented on a grey background screen, pointing either to the left, right, or both sides. Target sounds were monaural pure tones (carrier frequency between 512 and 575 Hz; 5 ms rise-time, 5 ms fall-time). In 25 % of the trials, a binaural distracting sound (300 ms duration) was played during the cue-target delay (50-650 ms range after cue offset). Trials with a distracting sound played from 50 ms to 350 ms after the cue offset were classified as DIS1; those with a distracting sound played from 350 ms to 650 ms after the cue offset were classified as DIS2; those with no distracting sound were classified as NoDIS. Please note that distracting sounds were uniformly paced in time between 50 and 650 ms after the cue offset. A total of 40 different ringing sounds were used as distracting sounds (clock-alarm, doorbell, phone ring, etc.) for each participant.

The cue and target categories were manipulated in the same proportion for trials with and without a distracting sound. In 25% of the trials, the cue was pointing left, and the target sound was played in the left ear, and in 25% of the trials, the cue was pointing right, and the target sound was played in the right ear, leading to a total of 50% of *informative* trials. In the other 50% of the trials, the cue was *uninformative*, pointing in both directions, and the target sound was played in the left (25%) or right (25%) ear. To compare brain responses to acoustically matched sounds, the same distracting sounds were played in each combination of cue category (informative, uninformative) and distractor condition (DIS1 or DIS2). Each distracting sound was thus played 4 times during the whole experiment, but no more than once during each single block to limit habituation.

Participants were instructed to categorize two target sounds as either high- or low-pitched sound, by either pulling or pushing a joystick. The target type (high or low) was manipulated in the same proportion in all conditions. The mapping between the targets (low or high) and the responses (pull or push) was counterbalanced across participants, but did not change across the blocks, for each participant. In order to account for the participants’ pitch-discrimination capacities, the pitch difference between the two target sounds was defined in a Discrimination Task (see below). Participants were informed that informative cues were 100 % predictive and that a distracting sound could be sometimes played. They were asked to allocate their attention to the cued side in the case of the informative cue, to ignore the distractors and to respond as quickly and correctly as possible. Participants had a 3.4 second response window. In the absence of the visual cue, a blue fixation cross was presented at the center of the screen. Subjects were instructed to keep their eyes fixated on the cross.

**Figure 1.**
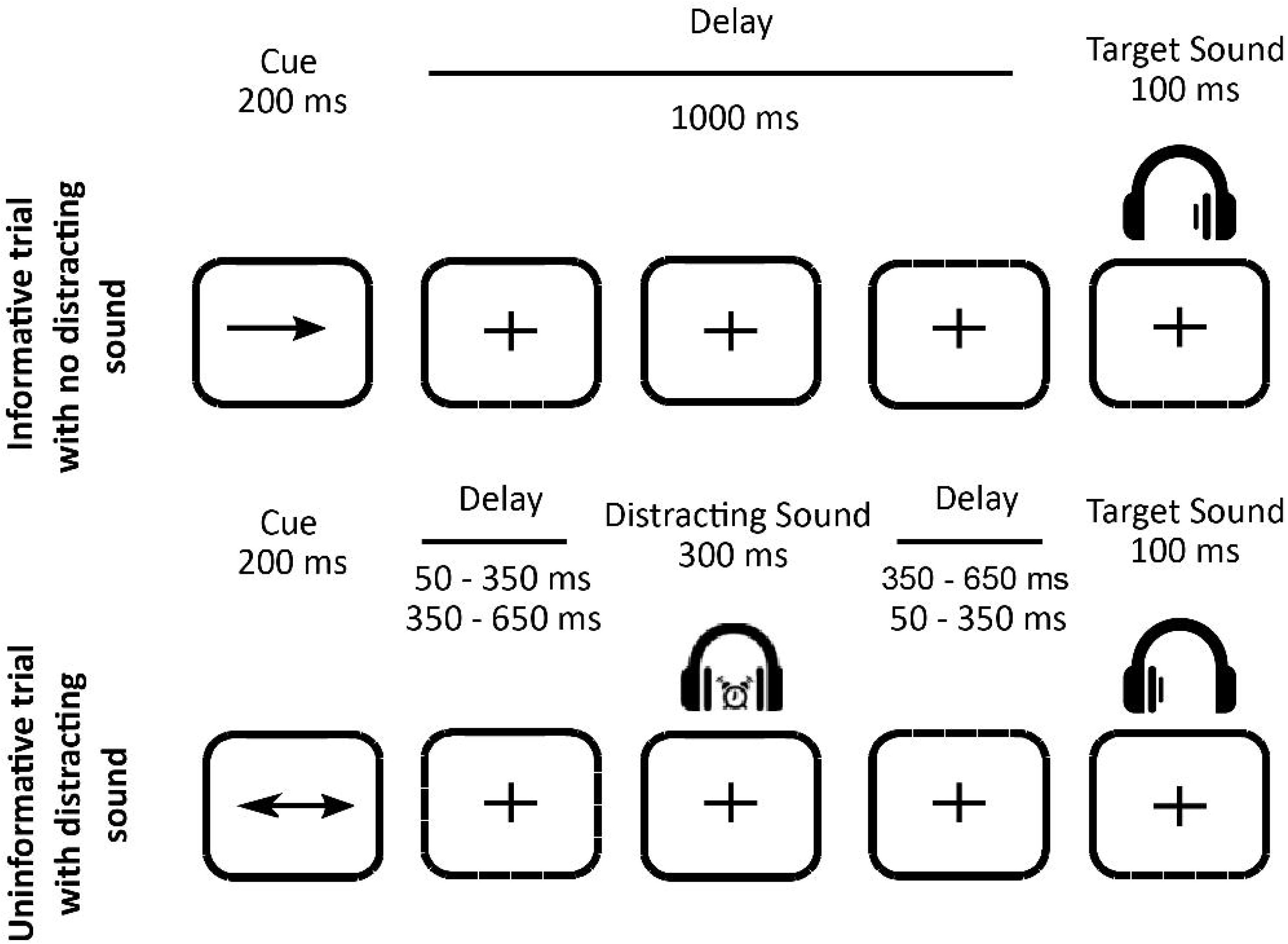
Protocol. Top row. Example of an informative trial with no distracting sound: a one-sided visual cue (200 ms duration) indicated in which ear (left or right) the target sound would be played (100 ms duration) after a fixed 1000-ms delay. Bottom row. Example of an uninformative trial with a distracting sound: a two-sided visual cue (200 ms duration) did not provide any indication in which ear (left or right) the target sound will be played. In 25 % of the trials, a binaural distracting sound (300 ms duration), such as a clock ring, was played during the delay between cue and target. The distracting sound could equiprobably onset in two different time periods after the cue offset: in the 50–350 ms range, or in the 350–650 ms range.

#### 2.2.2 Discrimination Task

Participants were randomly presented with one of the two target sounds: a low-pitched sound (512 Hz) and a high-pitched sound (575 Hz; two semitones higher), equiprobably in each ear (four trials per ear and per pitch). As described above, participants were asked to categorize the target sounds as either high- or low-pitched sound within 3 seconds.

#### 2.2.3 Procedure

Participants were seated in a sound-attenuated, magnetically shielded recording room, at a 50 cm distance from the screen. The response device was an index-operated joystick that participants moved either towards them (when instructed to pull) or away from them (when instructed to push). All stimuli were delivered using Presentation software (Neurobehavioral Systems, Albany, CA, USA). All sounds were presented through air-conducting tubes using Etymotic ER-3A foam earplugs (Etymotic Research, Inc., USA).

First, the auditory threshold was determined for the two target sounds differing by 2 semitones (512 and 575 Hz), for each ear, for each participant using the Bekesy tracking method (56). The target sounds were then presented at 25 dB sensation level (between 30 and 69.5 dB A in elderly, and between 37.5 and 50.75 dB A in young participants) while the distracting sounds were played at 55 dB sensation level (between 40 and 79.5 dB A in elderly, and between 47.5 and 60.75 dB A in young participants), above the target sound thresholds. Second, participants performed the discrimination task. If participants failed to respond correctly to more than 85% of the trials, the pitch of the high target sound was augmented, by half a semitone with a maximum difference of 3 semitones between the two targets (auditory thresholds were then measured with the new targets). Afterwards, participants were trained with a short sequence of the Competitive Attention Task. Finally, MEG and EEG were recorded while the subjects performed 10 blocks (64 trials each) leading to 240 trials in the NoDIS and 80 in the DIS conditions, for the informative and uninformative cues, separately. The whole session lasted around 80 minutes. After the MEG/EEG session, participants’ subjective reports regarding their strategies were collected.

### 2.3 Behavioral Data Analysis

For behavioral data analysis, a button press before target onset was considered as a false alarm (FA). A trial with no button-press after the target onset and before the next cue onset was considered as a miss trial. A trial with no FA and with a button-press after target onset was counted as correct if the pressed button matched the response mapped to the target sound, and as incorrect if otherwise. Reaction-times (RTs) to targets were analyzed in the correct trials only. To account for the heterogeneity of performances between-subjects and experimental conditions, we used linear mixed-effect (lme) models, as they allow for the correction of systematic variability (57) using the lme4 package (58) in R (59). By defining conditions as effects with random intercept and slope, we instructed the model to correct for any systematic differences in variability between the subjects (between-individual variability) and conditions (between-condition variability). To confirm the need for mixed nested models, we used a likelihood ratio analysis to test the model fit before and after sequential addition of random effects. We used the Akaike Information Criterion and the Bayesian Information Criterion as estimators of the quality of the statistical models generated (60). The influence of (1) age group (2 levels: young and elderly), (2) cue condition (2 levels: informative and uninformative), and (3) distractor condition (3 levels: NoDis, DIS1 and DIS2) on percentage of incorrect responses and median reaction times (RTs) of correct responses was tested. For post-hoc analysis, we used the Lsmean package (61) where p-values were considered as significant at p<0.05 and adjusted for the number of comparisons performed (Tukey method). Moreover, planned analyses of the CUE BENEFIT were carried out between groups on the differences in RTs Uninformative NoDIS – Informative NoDIS and of distractor effects on the differences in RTs NoDIS – DIS1 (as a measure of the AROUSAL BENEFIT) or DIS2 – DIS1 (as a measure of DISTRACTION COST), using unpaired t-tests (47, 48).

### 2.4 Brain Recordings

Simultaneous EEG and MEG data were recorded, although the data from the 7 EEG electrodes will not be presented here. The MEG data were acquired with a 275-sensor axial gradiometer system (CTF Systems Inc., Port Coquitlam, Canada) with a continuous sampling rate of 600Hz, a 0–150Hz filter bandwidth, and first-order spatial gradient noise cancellation. Moreover, eye-related movements were measured using vertical and horizontal EOG electrodes. The participant s head position relative to the gradiometer array was acquired continuously using coils positioned at three fiducial points; nasion, left and right pre-auricular points. The participant’s head position was checked at the beginning of each block to control head movements.

In addition to the MEG/EEG recordings, T1-weighted three-dimensional anatomical images were acquired for each participant using a 3T Siemens Magnetom whole-body scanner (Erlangen, Germany). These images were used for the reconstruction of individual head shapes to create the forward models for the source reconstruction procedures. The processing of these images was carried out using CTF’s software (CTF Systems Inc., Port Coquitlam, Canada). MEG data were pre-processed in the sensor space using the software package for electrophysiological analysis (ELAN pack; 48). Further analyses were performed using Fieldtrip (63), an open source toolbox for MATLAB and custom-written functions.

**Figure 2.**
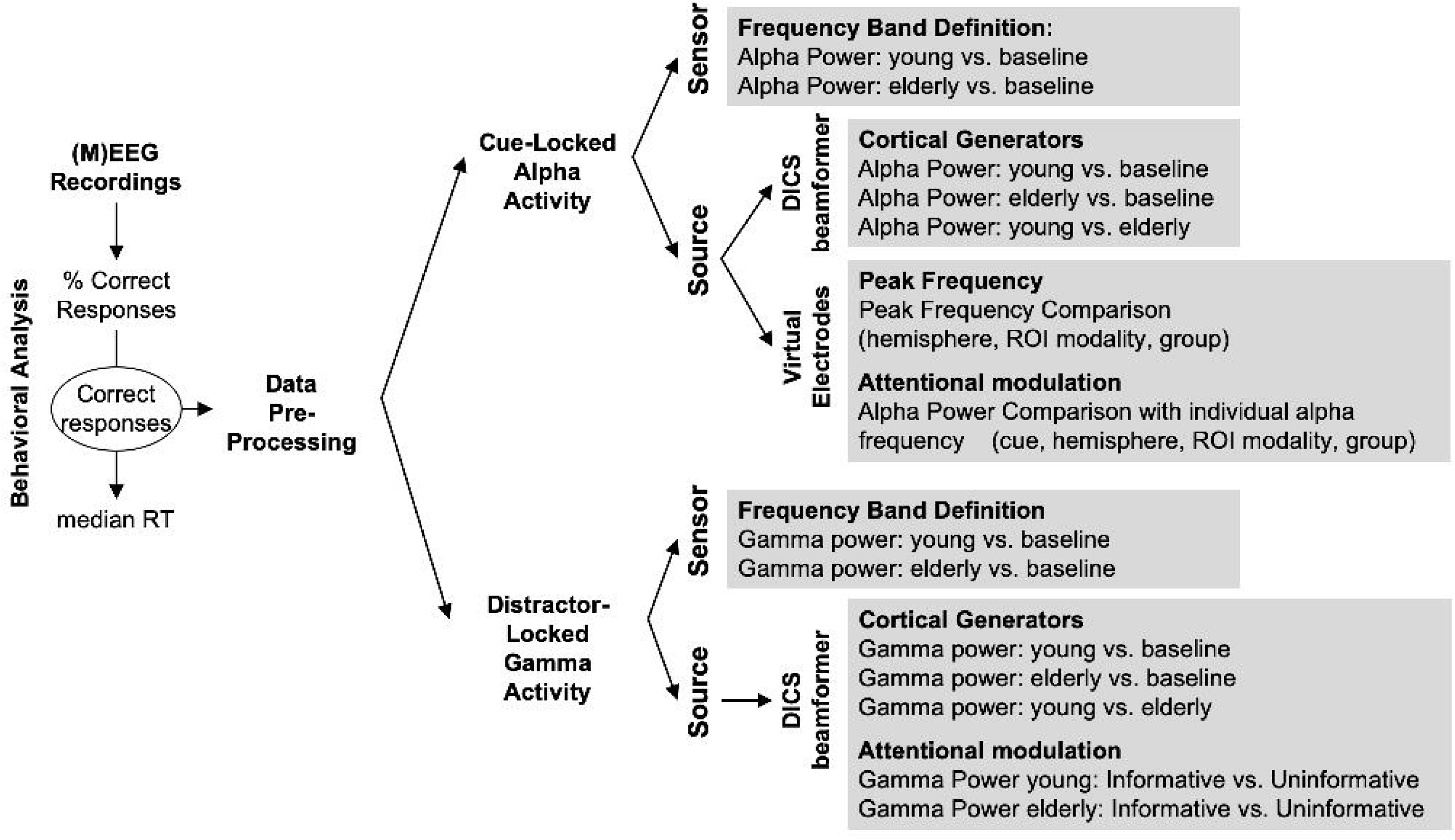
Outline of the analysis pipeline.

### 2.5 Data Pre-processing (Figure 2)

Only correct trials were considered for electrophysiological analyses. Data segments for which the head position differed for more than 10 mm from the median position during the 10 blocks were excluded. In addition, data segments contaminated with muscular activity or sensor jumps were excluded semi-manually using a threshold of 2200 and 10000 femtoTesla respectively. For all participants, more than 75 % of trials remained after rejection for further analyses. Independent component analysis was applied on the band-pass filtered (0.1-40Hz) data in order to remove eye-related (blink and saccades) and heart-related artifacts. Subsequently, components (four on average) were removed from the non-filtered data via the inverse ICA transformation. Data were further notch filtered at 50, 100 and 150Hz and high-pass filtered at 0.2Hz. Filtering was done using Butterworth filters of the third order.

Please note that for all subsequent source analyses, for each participant, an anatomically realistic single-shell headmodel (volume conduction model) based on the cortical surface was generated from individual head shape. A grid (source model) with 0.5-cm resolution was created using an MNI template, and then morphed (adjusted) into the brain volume of each participant using non-linear transformation. This procedure ensures that each grid-point would represent the same anatomical label across all subjects but with different subject-specific head positions.

### 2.6 Cue-Related Alpha Activity (Distractor-Free Trials)

#### 2.6.1 Alpha Band (Sensor level): Definition

This analysis was done to define the time-frequency range of alpha activity of interest. For all NoDIS (distractor free) trials, the oscillatory power (−3 to 3s relative to visual cue onset) was calculated using Morlet Wavelet decomposition with a width of four cycles per wavelet (m=7; 65) at center frequencies between 1 and 50 Hz, in steps of 1 Hz and 50 ms. For each group, activity of interest (between 0 and 1.2s post-cue and 5-45Hz) was contrasted to a baseline (−0.6 to −0.2s pre-cue) using nonparametric cluster-based permutation analysis (66).

#### 2.6.2 Alpha Band (Source level): Cortical Generators

To elucidate the possible brain regions underlying the sensor-level modulations, we have defined one post-cue (0.6 –1.0s) and one pre-cue (−0.6 to −0.2s) time-windows in two different frequency bands (9 and 13 ± 2 Hz). These time-frequency windows have been based on the results from the statistical contrasts in the sensor level (previous section) and from a previous study in an independent sample (6). For this purpose, we have used the frequency–domain adaptive spatial technique of dynamical imaging of coherent sources (DICS) (67). For each participant, using the data from all NoDIS trials, the cross-spectral density (CSD) matrix (−0.7 to 2 s, relative to cue onset, lambda 5%) was calculated using DPSS multitapers with a target frequency of 11 (±4) Hz. Leadfields for all grid points along with the CSD matrix were used to compute a common spatial filter that was used to estimate the spatial distribution of power for each time-frequency window of interest.

Afterwards, we performed the following analyses using nonparametric cluster-based permutation analysis (controlling for multiple comparisons in the source space dimension):

1. For each group and each alpha band separately, the post-cue time-frequency window was contrasted against the corresponding pre-cue baseline time-frequency window.
2. In order to investigate group differences in each alpha band, the baseline corrected power in the post-cue time-frequency window was compared between groups.

#### 2.6.3 Reconstruction of Time- and Frequency-Resolved Source Activity (Virtual Electrodes)

To resolve the time-frequency course of activity at the source level, the source space was subdivided into 69 anatomically defined brain parcels according to the Talairach Tournoux atlas (68, 69). ROIs of interest were defined based on the whole brain source analysis (previous section). Broadmann areas 17, 18 and 19 were defined as the visual regions of interest (ROIs); while areas 22, 41 and 42 were defined as the auditory ROIs, and areas 4 and 6 were defined as the motor ROIs, in each hemisphere. In order to reconstruct the activity at the source level, we computed the time-frequency signal of the ROIs defined above at the virtual electrode level. Using the linearly constrained minimum variance (LCMV) beamformer (70), spatial filters were constructed from the covariance matrix of the averaged single trials at sensor level (NoDIS trials, −0.8 – 2s, relative to cue onset, 1-20 Hz, lambda 5%) and the respective leadfield for all grid points. Spatial filters were multiplied by the sensor level data in order to obtain the time course activity at each voxel of interest within the defined ROIs.

Activity was averaged across all voxels within each ROI in each hemisphere. Thus, limiting our analysis to six ROIs (left/right auditory, left/right visual and left/right motor). For each ROI, the evoked potential (i.e., the signal averaged across all trials) was subtracted from each trial. Subsequently, the oscillatory power, of distractor-free trials, was calculated using Morlet Wavelet decomposition with a width of four cycles per wavelet (m=7; 65) at center frequencies between 5 and 20 Hz, in steps of 1 Hz and 50 ms (see Sup Figure 1).

#### 2.6.4 Impact of Age on Alpha Peak Frequency

For all NoDIS (distractor free) trials, alpha power (computed using Morlet Wavelets) was averaged between 0.6 and 1s for each ROI, to extract the power spectrum in each subject. Afterwards, individual alpha peak frequency (iAPF) was defined separately for each ROI, in each subject. For the auditory and motor ROIs, the peak was defined as the frequency with the maximum alpha power decrease relative to the baseline (−0.6 to −0.2s pre-cue onset) between 5 and 15 Hz. For the visual virtual electrodes, the peak was defined as the frequency with the maximum alpha power increase relative to the baseline.

The iAPFs were fit into a lme model with the following factors: (1) age group (2 levels: young and elderly), (2) ROI modality (3 levels: auditory, visual and motor), and (3) hemisphere (2 levels: left and right). Fixed and random effects were chosen following the same procedure as for behavioral data analysis (part 2.3). For post-hoc analysis, we used the Lsmean package, similar to previous analysis.

#### 2.6.5 Impact of Age and Top-down Attention on Alpha Power

In order to investigate the impact of ageing and top-down attention on alpha power at the source level, for each participant, baseline-corrected (relative to −0.6 to −0.2s pre-cue) alpha power (computed using Morlet Wavelets) was computed at the iAPF of each ROI. Power values were fit into a lme model with the following factors: (1) age group (2 levels: young and elderly), (2) ROI modality (3 levels: auditory, visual and motor), (3) hemisphere (2 levels: left and right), and (4) cue condition (2 levels: informative and uninformative). Fixed and random effects were chosen following the same procedure as for behavioral data analysis (part 2.3). Post-hoc tests were also computed in a similar manner to the aforementioned part (2.3).

### 2.7 Distractor-Related Gamma Activity

#### 2.7.1 Gamma Band (Sensor level): Definition

This analysis was done to define the time-frequency range of gamma activity of interest. First, for each distractor onset time-range, we have created triggers for surrogate (fake) distractors in the NoDIS trials with similar distribution over time to the real distractors i.e. in the NoDIS trials we have added “fake” triggers around which the surrogate or “fake” trials would be epoched. These trials would serve as a baseline for trials with “real” distractors by being subtracted. Afterwards, the oscillatory power, of distractor and (surrogate distractor) trials was calculated using Morlet Wavelet decomposition with a width of four cycles per wavelet (m=7; 65) at center frequencies between 40 and 150 Hz, in steps of 1 Hz and 10 ms. For each group, activity of interest (defined between 0 and 0.35s post-distractor onset and 50-110Hz) was contrasted between distractor and surrogate trials using a nonparametric cluster-based permutation analysis (66). This contrast extracts distractor-related activity clear of cue-related activity.

#### 2.7.2 Gamma Band (Source level): Cortical Generators

In order to estimate the brain regions driving the sensor-level distractor-related gamma activity (0.1-0.3 s post-distractor onset, 60 to 100Hz), we have utilized the frequency–domain (DICS) (67), similar to analysis for the alpha band. Data, from both surrogate and real distractors were concatenated, and the cross-spectral density (CSD) matrix (−0.2 to 0.6 s, relative to real/surrogate distractor onset, lambda 5%) were calculated using DPSS multitapers with a target frequency of 80 (±30) Hz. Leadfields for all grid points along with the CSD matrix were used to compute a common spatial filter that was used to estimate the spatial distribution of power for the time-frequency windows of interest.

For each participant, we estimated source-level activity (0.1-0.3 s post-distractor onset, 60 to 100Hz) for both cue categories concatenated. We performed three analyses:

1. To characterize the brain areas activated in the gamma band during the distracting sound presentation, for each group distractor-related gamma activity was contrasted to surrogate distractor-related gamma activity.
2. To investigate group differences in the processing of the distracting sound, surrogate-corrected gamma activity (surrogate distractor-related gamma activity was subtracted from distractor-related gamma activity) was compared between groups.
3. To investigate the impact of top-down attention on gamma activity, for each group separately, surrogate-corrected gamma activity in the informative cue condition was contrasted to surrogate-corrected gamma activity in the uninformative cue condition.

All tests have been carried out using non-parametric cluster-based permutation analysis (66). Please note, that for these tests, cluster-based permutations control for multiple comparisons in the source space dimension.

## 3 Results

### 3.1 Behavioral Analysis

Participants correctly performed the discrimination task in 95.37 ± 0.29 SEM % of the trials. The remaining trials were either incorrect trials (4.62 ± 0.29 SEM %), missed trials (0.49 ± 0.09 %) or trials with FAs (0.14 ± 0.03 %). In order to investigate behavioral performances (percentage of incorrect responses and RTs), lme models with 3 factors: (1) age group (2 levels: young and elderly), (2) cue condition (2 levels: informative and uninformative), and (3) distractor condition (3 levels: NoDis, DIS1 and DIS2) were used.

#### 3.1.1 Behavioral Analysis: Incorrect Response Percentage (Figure 3)

The best lme model accounted for the interindividual variability (subject as intercept). Only a significant main effect of the distractor condition (F(2, 52) = 8.9, p < 0.01, η^2^ = 0.21) was found on the percentage of incorrect responses, with no main effect of group (F(1, 15) = 2.1, p = 0.94, η^2^ = 0.07). Post-hoc tests indicated that participants, from both groups, committed more errors in the late DIS2 condition in comparison to the early DIS1 (p < 0.01) and the NoDIS (p < 0.001) conditions.

**Figure 3.**
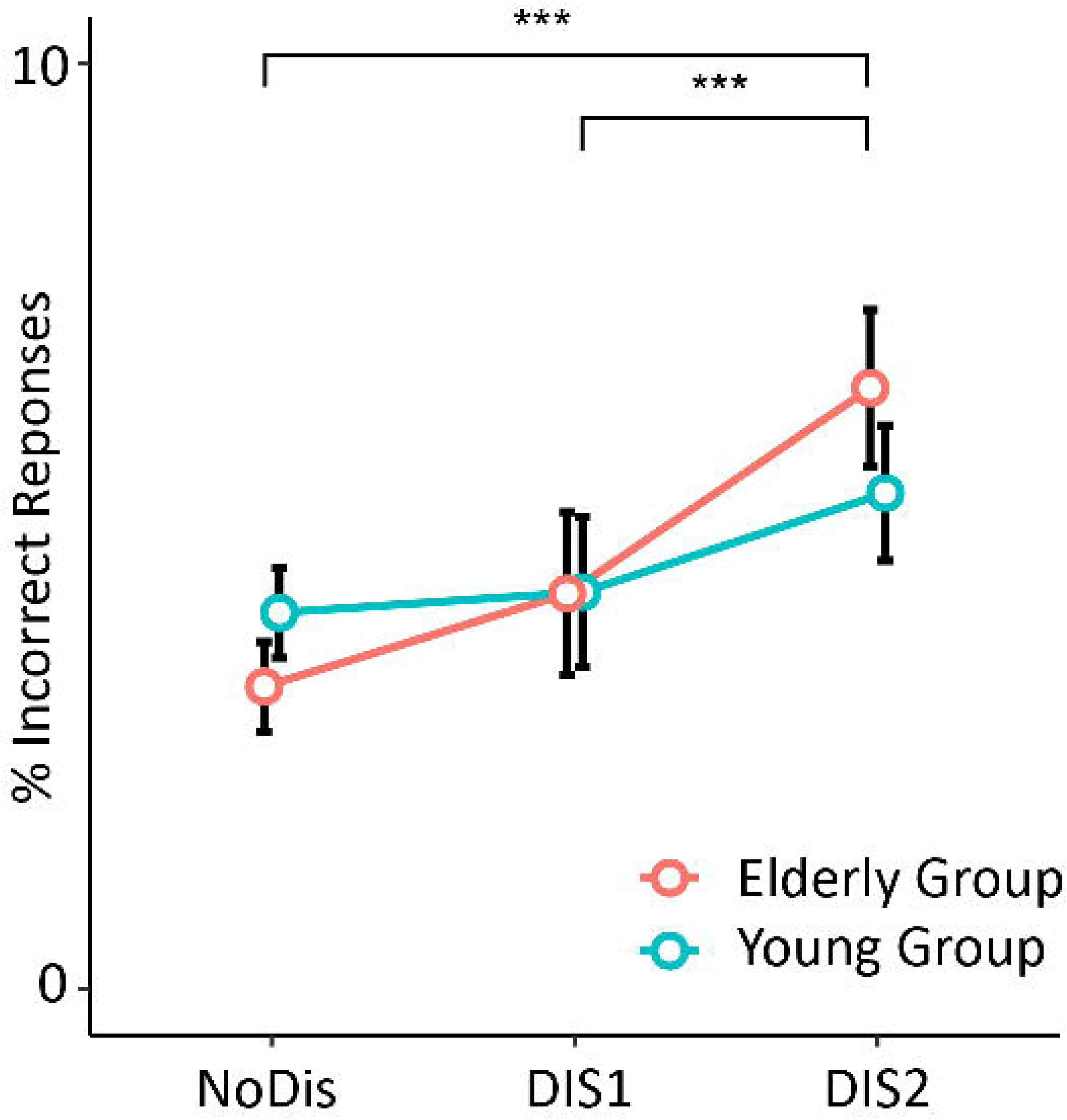
Percentage of Incorrect Responses averaged across cue conditions, according to distractor conditions, for both groups. *** *P* < 0.001. Error bars represent SEM.

#### 3.1.2 Behavioral Analysis: Median Reaction Times (Figure 4)

The best model accounted not only for the interindividual variability (subject as intercept) but also for the heterogeneity of the distractor effect (slope) across subjects. A main effect of the cue category (F(1, 26) = 4.69, p = 0.03, η^2^ = 0.22) was found on reaction times. Participants were faster when the cue was informative in comparison to the uninformative cue. In addition, we found a main effect of the distractor condition (F(2, 52) = 37.04, p < 0.01, η^2^ = 0.64). Post-hoc tests indicated that in comparison to the NoDIS condition, participants were faster in the early DIS1 condition (p = 0.001) but slower in the late DIS2 condition (p < 0.001). In addition, participants were faster in the early DIS1 than in the late DIS2 condition (p < 0.01). Most interestingly, we found a significant interaction between age and the distractor factors (F(2, 52) = 3.77, p = 0.036, η^2^ = 0.17). Post-hoc tests indicated that elderly participants tended to be slower than the young ones in the late DIS2 condition (p = 0.069); while no significant group difference was found in NoDIS and DIS1 conditions (p = 0.21 and p = 0.28, respectively).

Planned unpaired t-tests between groups confirmed that the distraction cost (DIS2 – DIS1) was significantly more pronounced for the elderly group (p < 0.01); whereas the arousal benefit (NoDIS – DIS1; p = 0.29) and the cue benefit (Uninformative NoDIS – Informative NoDIS; p = 0.83) were not significantly different between groups.

**Figure 4.**
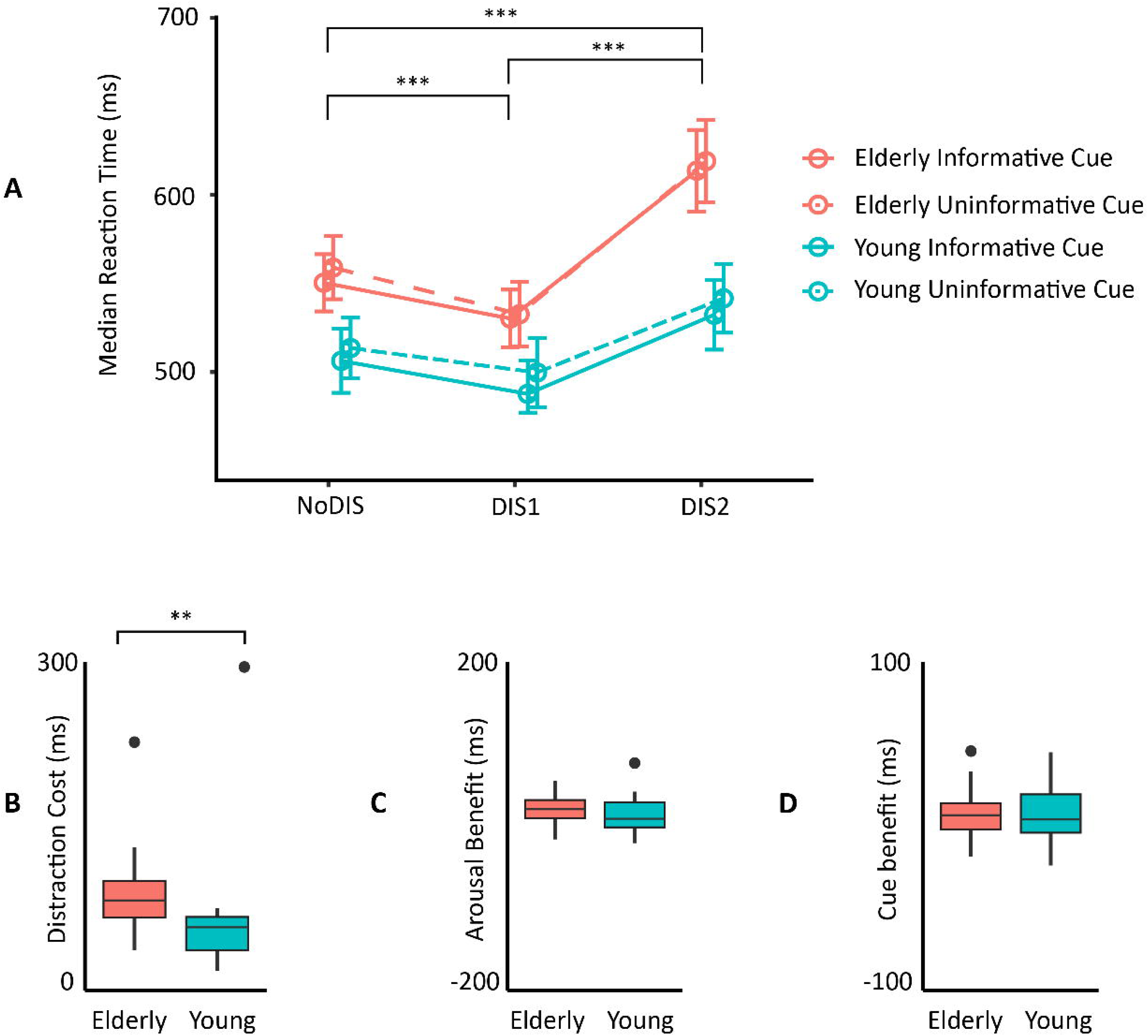
(A) Median RTs in each group according to cue information and distractor conditions. Error bars represent SEM. Boxplots of the Distraction cost (B), the Arousal benefit (C), and the Cue benefit (D), for each group. Within each boxplot (Tukey method), the horizontal line represents the median, the box delineates the area between the first and third quartiles (interquartile range). ** *P* < 0.01.

### 3.2 Cue-related Alpha Activity

#### 3.2.1 Alpha Band (Sensor level): Definition

As shown in figure 5, upon contrasting post-cue activity to baseline activity, the younger displayed a negative cluster (p < 0.001), indicating a decrease in alpha power (desynchronization), relative to the baseline, between 200 and 1200 ms and in lower alpha frequencies (7-11Hz). Spatially, this decrease was widespread during cue processing (early period: 200 and 600ms) but became centered around left temporo-parietal sensor in the late period (600 – 1000ms). This late alpha desynchronization was accompanied by a trending positive cluster indicating an increase in alpha power (synchronization; p = 0.084), relative to the baseline, in higher alpha frequencies (11-15Hz) centered on right occipito-temporal sensors.

In comparison, the elderly group displayed only one negative cluster (p < 0.001), indicating an alpha desynchronization between 200 and 1200 ms in alpha frequencies; but no significant alpha synchronization was found.

**Figure 5.**
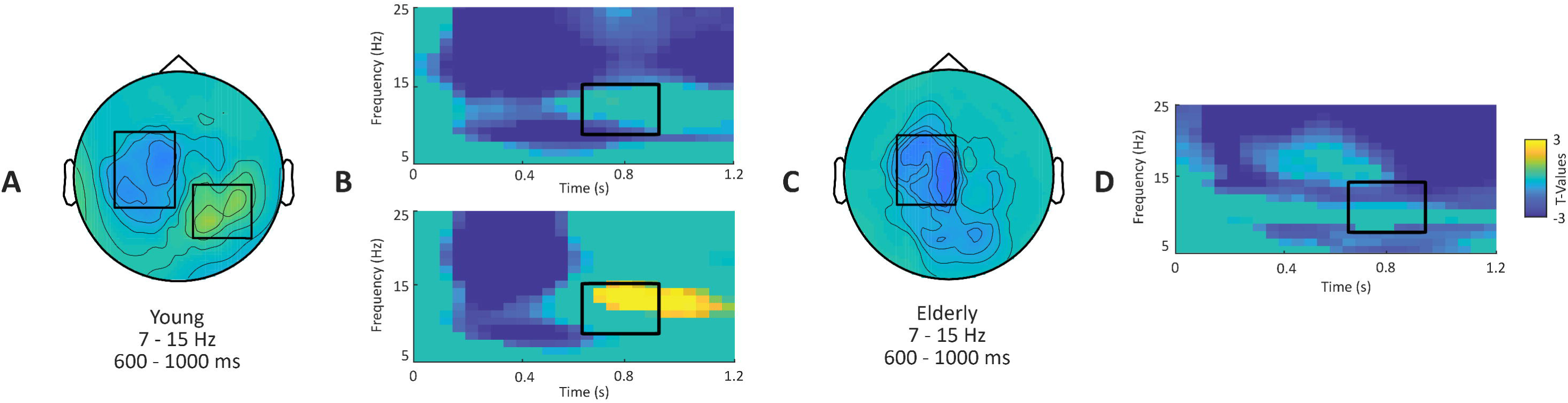
**Sensor level alpha activity.** Comparison between post-cue activity of interest (between 0 and 1.2s post-cue and 5-45Hz) to baseline (−0.6 to −0.2s pre-cue) for young and elderly participants. **A-C.** Topographical maps represent T-values, masked at p < 0.1, for the comparison in the alpha band (7-15 Hz and 0.6-1s post-cue). **B-D.** Time-frequency representations depict T-values, of the same tests, masked at p<0.1 and averaged across sensors highlighted by black boxes on the corresponding topographical maps.

#### 3.2.2 Alpha Band (Source level): Cortical Generators

Based upon the sensor level results and using DICS beamformer sources, for each group, post-cue (0.6-1s) low (7-11Hz) and high (11-15Hz) alpha source activities were contrasted against the pre-cue baseline (−0.6 to −0.2s) (see Figure 6 A & B).

The aim of this test was to highlight the brain regions that are activated in each group in the low and high alpha bands. In the young participants, we found a negative cluster (compared to baseline; p = 0.002), indicating a decrease in low alpha power in auditory cortices and in fronto-parietal regions with a maximum around the left motor cortices. We also found a positive cluster (compared to baseline; p = 0.003), indicating an increase in high alpha power in occipital areas. These findings in the young participants are in agreement with our previous findings in another group of young adults (71). In the elderly participants, we found a negative cluster (compared to baseline; p < 0.001), similar to the one in young participants in the low alpha band. However, in the high alpha band, we found no positive occipital cluster, but a negative one (p < 0.001) indicating a power decrease in frontal, motor and auditory areas.

**Figure 6.**
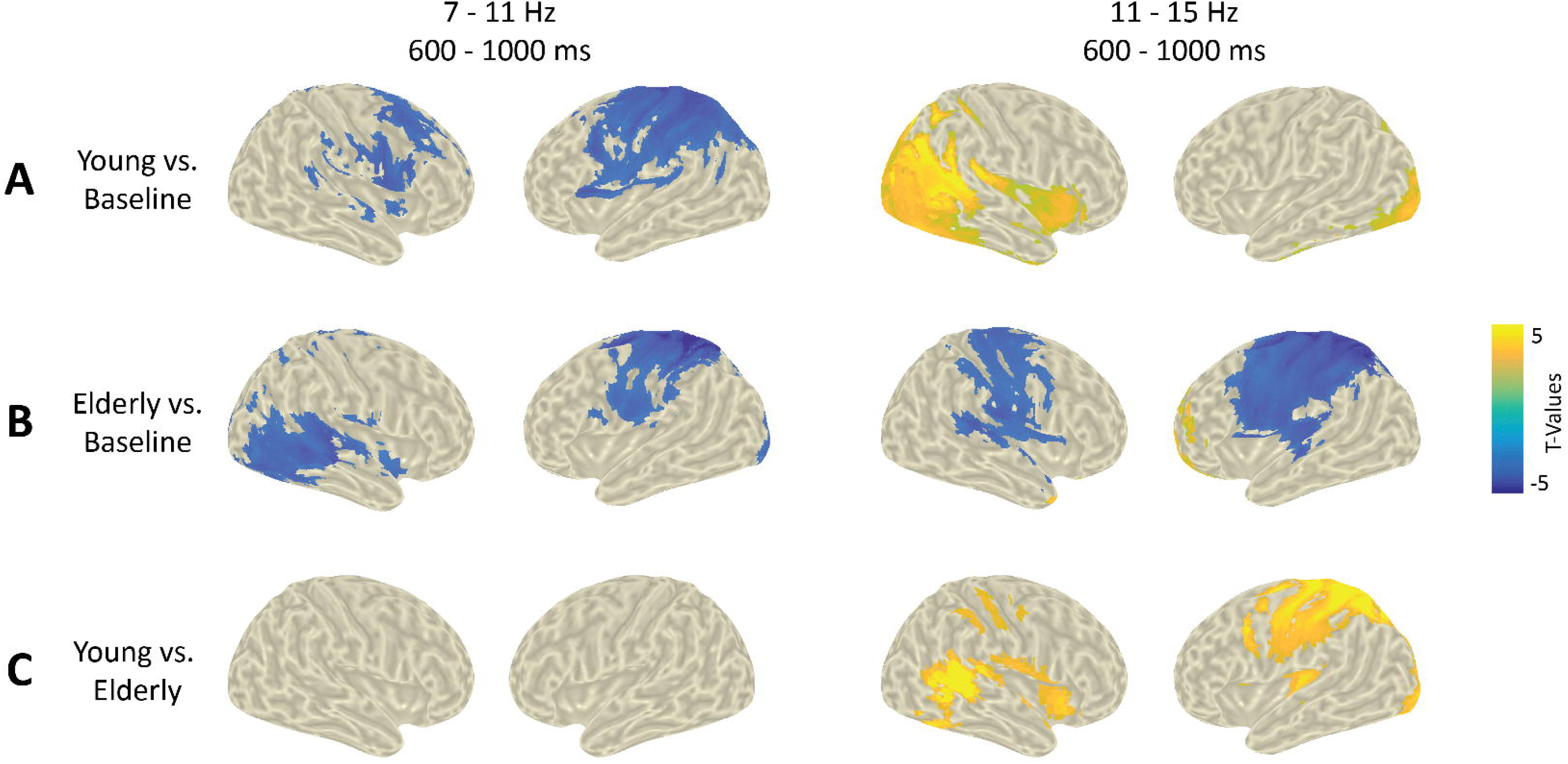
**Source level alpha activity. A-B.** Distributions of T-values, masked at *p* < 0.05, from cluster-based permutation tests contrasting time-frequency windows of interest (low and high alpha) against baseline activity at the source level for each group. **C**. Distributions of T-values, masked at *p* < 0.05, from cluster-based permutation tests contrasting baseline-corrected time-frequency window of interest between groups. Yellowish/blueish colors indicate a greater activity in the young/elderly group, respectively.

#### 3.2.3 Alpha Band (Source level): Group Comparison

The aim of this test was to highlight the brain regions that are differentially activated in the elderly and young groups, in the alpha bands. Baseline corrected alpha power in the low and high alpha bands was contrasted between groups using non-parametric cluster-based permutation testing (Figure 6C). No significant group differences were found in the low alpha band. A significant positive cluster (p < 0.001) was found, indicating greater power in the high alpha in young compared to elderly participants in the auditory cortices, in occipital regions and in the left motor cortices.

#### 3.2.4 Alpha band (Source level): Alpha peak frequency

In order to investigate the alpha peak frequency in virtual ROIs, lme models with 3 factors: (1) age (2 levels: young and elderly), (2) ROI modality (3 levels: auditory, visual and motor), and hemisphere (2 levels: left and right) were used. The best model accounted for interindividual variability (subject as intercept). This lme model yielded a significant interaction between age and ROI modality (F(2,52) = 4.1, p = 0.018, η^2^ = 0.11). Post-hoc tests indicated that, only in the young group, the alpha peak frequency was significantly higher in the visual ROI (mean = 12.32 Hz ± 0.44 SEM) than in the auditory (mean = 9.39 Hz ± 0.58 SEM) and the motor ROIs (mean = 10 Hz ± 0.58 SEM) (p < 0.001 and p = 0.01, respectively), as shown in Figure 7.

**Figure 7.**
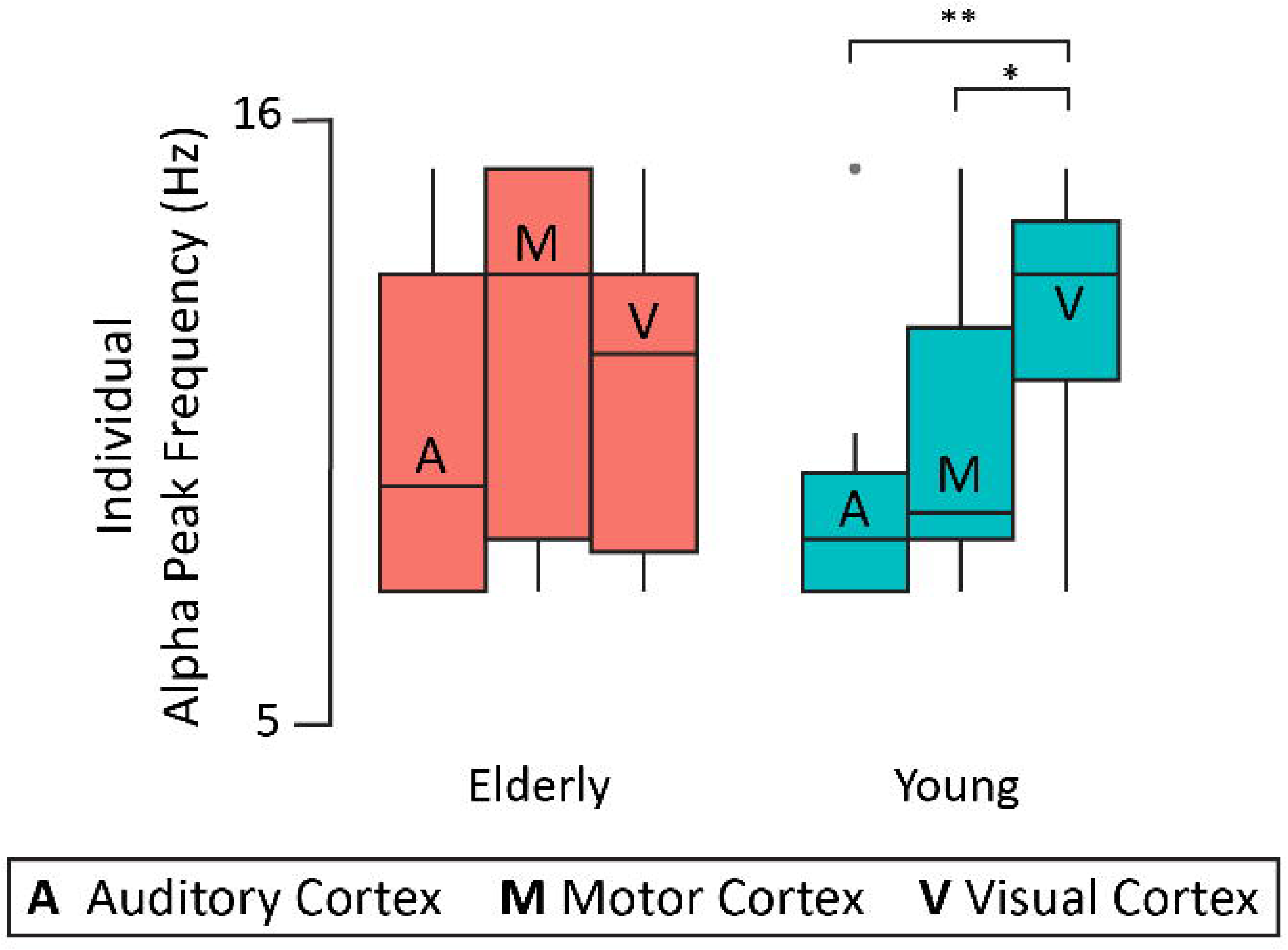
Individual alpha peak frequency in virtual ROIs (averaged across hemispheres) for each group. * p < 0.05, ** p < 0.01. Within each boxplot (Tukey method), the horizontal line represents the median of iAPF, the box delineates the area between the first and third quartiles (interquartile range).

#### 3.2.5 Alpha band (Source level): Age and Top-down Modulation

We investigated the impact of age and of cue-related top-down attention on alpha activity, at the individual alpha peak frequency, in the virtual ROIs. Lme models were used with 3 factors: (1) age (2 levels: young and elderly), (2) modality (3 levels: auditory, visual and motor), hemisphere (2 levels: left and right), and cue condition (2 levels: informative and uninformative). The best model accounted not only for the large interindividual variability (subject as intercept) but also for the heterogeneity of the hemisphere and modality effects (slope) across subjects. This lme model yielded several significant main effects and interactions (listed in Table 2).

The first highest-order significant interaction of interest was the three-level interaction between group, cue condition and ROI modality (F(2,54) = 4.54, p = 0.01, η^2^ = 0.02, see Figure 8). Post hoc testing revealed that for the young group, in the auditory and motor ROIs alpha power was significantly lower in the informative condition compared to the uninformative one (p < 0.001), while there was a trend for a larger alpha power in the informative condition compared to the uninformative one in the visual ROI (p = 0.08). For the elderly group, we found significantly lower alpha power in the informative condition compared to the uninformative one in the auditory, motor and visual ROIs (p = 0.0027, p = 0.0023 and p = 0.034, respectively).

Post hoc testing of the two-level interaction between group and modality (F(2,54) = 4.11, p = 0.03, η^2^ = 0.04) revealed that the visual alpha power increase (synchronization) was significantly higher for the young group (p = 0.015); while there was a trend for a more prominent motor alpha power decrease (desynchronization) for the elderly group (p = 0.086); with no significant difference between groups for the auditory desynchronization (p = 0.59). These effects are also visible on the time courses of alpha power in each ROI (Sup Figure 2).

Post hoc testing of the two-level interaction between hemisphere and modality (F(2,54) = 13.6, p <0.001, η^2^ = 0.013) revealed that the alpha power decrease (desynchronization) was significantly more prominent for the left than the right hemisphere in the motor ROIs (p < 0.001), with trends in the visual (p = 0.05) and auditory (p = 0.06) ROIs (see Sup Figure 2).

**Table 2.** Significant results of the lme model testing the modulation of alpha activity by group, cue condition, ROI modality and hemisphere.

| ***EFFECT*** | ***F*** | ***P*** |
| --- | --- | --- |
| **Group** | 5.36 | 0.03 |
| **Cue Condition** | 37.57 | $< 0.001$ |
| **Hemisphere** | 15.3 | $< 0.001$ |
| **Modality** | 53.22 | $< 0.001$ |
| **Group X ROI Modality** | 4.11 | 0.03 |
| **Cue Condition X ROI Modality** | 8.26 | $< 0.001$ |
| **Hemisphere X ROI Modality** | 13.62 | $< 0.001$ |
| **Group x Cue Condition x ROI Modality** | 4.54 | 0.01 |

**Figure 8.**
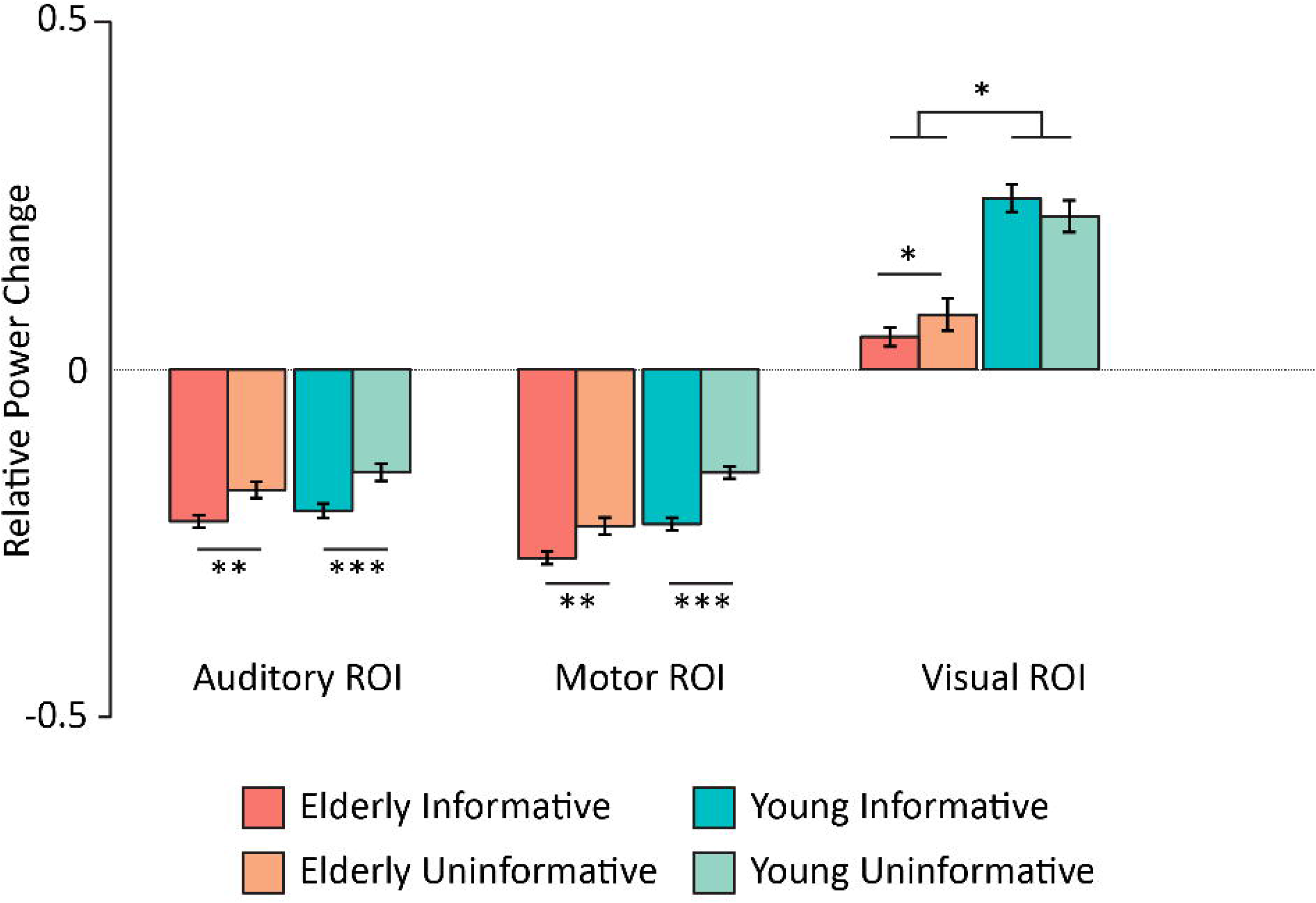
Mean alpha power (relative to baseline) centered around the alpha-peak frequency for each region and each participant, averaged between 600 and 1000 ms (post-cue onset) for each cue condition. *P < 0.05, **P < 0.01, ***P < 0.001. Error bars represent SEM.

### 3.3 Distractor-Related Gamma Activity

#### 3.3.1 Gamma-band (Sensor level): Definition

For each group, real-distractor high-frequency activity was contrasted to surrogate-distractor activity using non-parametric cluster-based permutation testing. As shown in Figure 9, for each group, this contrast revealed a significant positive cluster (young: p < 0.001, elderly: p = 0.002), indicating an increase in gamma activity centered spatially around left and right temporal sensors, temporally between 0.1 and 0.3s post-distractor onset, and frequency-wise between 60 and 100 Hz.

**Figure 9.**
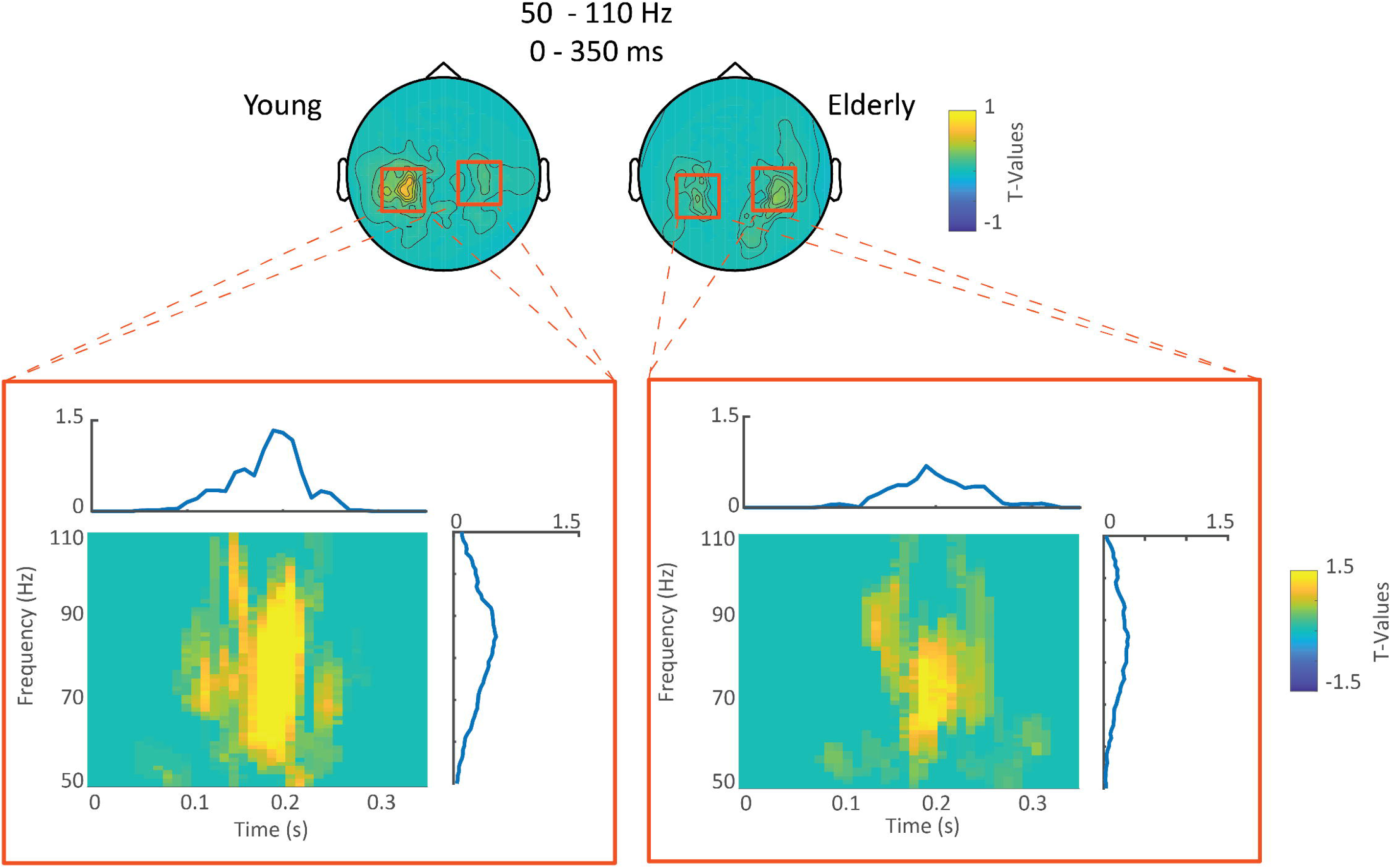
**Sensor level gamma activity.** Upper row: Topographical maps averaged between 50-110 Hz and 0-0.35 seconds post-distractor onset, of the T-values, masked at p < 0.05 of the contrast between distractor and surrogate distractor gamma activity for each group. Lower row: Time-frequency representations of T-values (of the aforementioned test) of the sensors highlighted by red boxes. Note that each TF plot is surrounded by the frequency distribution of T-values of the aforementioned sensors averaged across the time dimension and by the time distribution of T-values of the aforementioned sensors averaged across the frequency dimension.

#### 3.3.2 Gamma Band (Source level): Cortical Generators

Based upon sensor level results, we have computed DICS beamformer sources for each participant between 60-100Hz and 0.1-0.3s post-distractor. Once again, for each group, real-distractor gamma activity was contrasted to that of surrogate distractors using non-parametric cluster-based permutation testing.

The aim of this test was to highlight the brain regions that are activated in each group in the gamma band. For each group, a significant positive cluster (p < 0.01) was found, indicating a bilateral increase in gamma activity notably in the auditory cortices, the temporo-parietal junctions and the ventro-lateral prefrontal cortices (Figure 10A & B). Other regions included the middle and posterior cingulate gyri, pre- and post-central gyri, the precuneus, and the inferior temporal gyri.

#### 3.3.3 Gamma Band (Source level): Group Comparison

The aim of this test was to highlight the brain regions that were differentially activated in the elderly and young groups, in the gamma band. For each participant, real distractor source-level data (60-100 Hz, 0.1-0.3s) were corrected by subtracting surrogate-distractor activity. Corrected distractor gamma activity was contrasted between groups using non-parametric cluster-based permutation testing. A significant positive cluster (p = 0.01) was found notably in the left dorsolateral prefrontal cortices (Figure 10C). Other regions included the left post-central gyrus and the left supplementary motor area. All these regions displayed a higher gamma activation for the young group compared to the elderly group.

**Figure 10.**
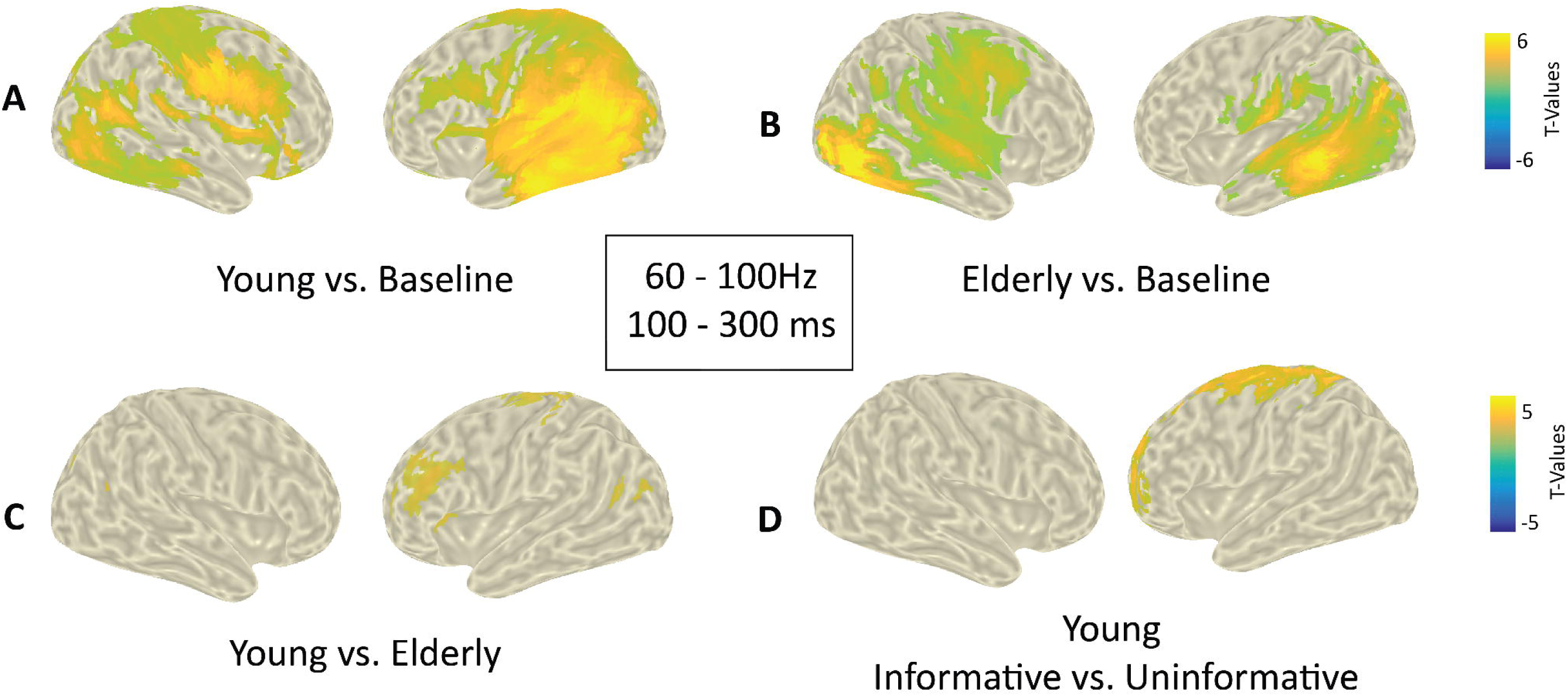
**Source level gamma activity. A&B**. Distributions of T-values, masked at p<0.05, from Cluster Based Permutation tests contrasting real and surrogate distractor gamma activity (60-100Hz and 0.1-0.3 post-distractor) for elderly and young groups, respectively, at the source level. **C**. Between group comparison of surrogate-corrected distractor gamma activity (60-100Hz and 0.1-0.3 post-distractor) at the source level. Positive T-values indicate stronger gamma activity in the young group. Yellowish/blueish colors indicate a greater activity in the young/elderly group, respectively. **D**. Between cue condition comparison of surrogate-corrected gamma activity to distractor in the young group. Positive T-values indicate stronger gamma activity in the informative cue condition. Yellowish/blueish colors indicate a greater activity in the informative/uninformative condition, respectively. Note that no significant effect was found in the elderly group.

#### 3.3.4 Gamma Band (Source level): Top-down modulation

The aim of this test was to highlight the brain regions that are modulated by top-down attention, i.e. differentially activated in the informative and uninformative cue conditions, in the gamma band, in each group. For each participant, the real distractor source-level data (60-100 Hz, 0.1-0.3s) were baseline corrected by subtracting surrogate-distractor activity. For each group separately^1^, corrected distractor gamma activity was contrasted between the two cue categories (informative vs. uninformative) using non-parametric cluster-based permutation testing. Only for the young group, a significant positive cluster (p = 0.02) was found in the dorso- and ventromedial prefrontal cortices, the anterior cingulate gyrus, the pre- and post- central gyri, the supplementary motor area, the superior parietal lobule, in the left hemisphere (Figure 10D). All regions displayed a significantly higher gamma activation for trials when the distracting sound was preceded by an informative cue rather than an uninformative cue.

## 4 Discussion

The present results demonstrate that ageing differently impacts TD (anticipatory) and BU (capture) attentional processes. Behavioral measures of TD attention seemed unchanged while distractibility was exacerbated with ageing. Electrophysiologically, during the anticipation of visually cued sounds, the alpha power decrease in the relevant auditory cortices was preserved; whereas the alpha increase in the irrelevant visual areas was reduced, in elderly compared to young adults. Moreover, in response to distracting sounds, similar activation of the ventral BU attentional network, but reduced activation of lateral prefrontal regions was observed in the gamma band, with ageing.

### 4.1 Impact of Ageing on Behavioral Measures of Attention

Behaviorally, both groups benefited from the cue information to faster discriminate the target pitch. This is in line with previous studies suggesting that TD attentional orienting is not affected by ageing (72–75). In trials with distracting sounds, both groups displayed a similar reaction time (RT) pattern: in agreement with previous studies (35, 46 47), participants were faster after early distracting sounds and slower after late distracting sounds, than with no distractors. This pattern could be explained in light of the effects triggered by distracting sounds (review in 47,76): (1) a persistent increase in arousal resulting in a RT reduction and (2) a strong transient attentional capture (orienting) effect associated with a RT augmentation. The behavioral net effect of the distracting sound varies according to the time interval between the distracting and the target sounds (see Supplementary Figure 4). Thus, with the present paradigm, the difference in RT to targets preceded by late distracting sounds (DIS2) and early distracting sounds (DIS1) provides a good approximation of the distraction effect with little contamination by arousal increases. Importantly, in comparison to the young group, elderly participants displayed a larger distraction effect. This increased susceptibility to task-irrelevant distractors is a recurrent finding in the literature using unimodal (visual or auditory) or cross-modal paradigms (77–81).

### 4.2 Impact of Ageing on TD Attentional Mechanisms

In the present study, we replicate with a slightly different protocol previous findings in an independent sample of young adults. We found, in comparison to the baseline, during the anticipation of a visually-cued auditory target, (1) an alpha decrease around 9Hz in task-relevant auditory regions accompanied by (2) an increase in alpha power around 13 Hz in task-irrelevant visual regions (6). According to the dominant hypotheses on the functional role of alpha oscillations (22,25,26), reduced/enhanced alpha power reflect increased/decreased cortical excitability in task relevant/irrelevant areas, respectively. In the light of previous fMRI studies demonstrating how TD attention enhances/reduces activity in the sensory cortex of the attended/unattended modality (e.g. 82), our results suggest that alpha desynchronization and synchronization result from facilitatory and suppressive mechanisms of TD attention, respectively, augmenting and reducing cortical excitability.

Noteworthy, when taking into account interindividual variability in alpha peak frequency, we observed that elderly participants and young adults, seem to display a similar alpha desynchronization in the task-relevant auditory cortices and to similarly modulate this alpha decrease by top-down attention according to the cue information. However, elderly participants present a reduced alpha synchronization in the task-irrelevant visual cortices. These findings suggest that with ageing, at least during anticipation of task-relevant stimuli, suppressive TD attentional mechanisms become defect, while facilitatory mechanisms would be preserved, in line with (a) previous studies of alpha oscillations during visual attention (44, 46), and (b) previous fMRI studies showing that filtering out task-irrelevant information is reduced with ageing (e.g. 78). Moreover, we found a larger alpha desynchronization in the motor areas in the elderly, in agreement with an ageing-related stronger engagement of motor regions during response preparation (e.g. 41,83). The elderly could rely more on motor preparation processes as a compensatory mechanism to their reduced attention filtering of irrelevant information, resulting in comparable performances to younger adults at the behavioral level.

Finally, a novel finding in the present study is the decreased differentiation between alpha peak frequency of different sensory regions, with ageing. This finding is in line with studies showing changes in alpha peak frequency across brain regions, through the life-span (e.g. 84,85). Yet, this raises the question: is this decreased differentiation causal to the failure of older participants to filter out irrelevant visual information? Or does the alteration of suppressive attentional mechanisms result in a reduced alpha frequency specificity in the ageing brain? These questions remain for future investigations to answer.

### 4.3 Impact of Ageing on Distractor processing

To the best of our knowledge, this is the first investigation of gamma activity to highlight the impact of ageing on auditory BU attentional processes. In both groups, gamma activation in bilateral auditory cortices, and several other brain regions including the temporo-parietal junctions and the right ventro-lateral prefrontal cortex, was observed in response to an unexpected salient distracting sound. These regions are part of the well-established ventral BU attentional network observed in fMRI (e.g. 86–89) and electrophysiological (e.g. 36,90) studies. In the fMRI literature, it is rather unclear how ageing impacts the BU network with evidence of preserved (91), reduced (92) and even augmented (93) activation with ageing. The present results suggest a similar activation of the ventral BU attentional network in the gamma band in elderly and young adults.

Importantly, younger participants demonstrated higher gamma activation than elderly ones in regions outside the ventral network: mainly in the dorsolateral prefrontal cortices (PFC) in the left hemisphere. In addition, only young participants were found to modulate gamma activity by TD attention according to the preceding cue information. In agreement with previous results (36), gamma activity was larger after an informative cue in comparison to an uninformative cue in the medial PFC.

This observed decline in PFC activation, in particular in its lateral part, in the elderly is in agreement with structural alterations in the prefrontal cortices with ageing (e.g. 94). The lateral PFC has been proposed to be part of the TD inhibitory control system: In the non-human primate, Suzuki and Gottleib (14) demonstrated that the inactivation of the dorsolateral PFC results in impaired distractor suppression, i.e. increased distractibility. In humans, reduced activity in the lateral PFC caused by ageing or strokes has been linked to enhanced distractor processing (e.g. 95,96); while stimulation of the lateral PFC by utilizing transcranial direct-current stimulation (tDCS) decreased the behavioral cost of distractors (97). Finally, increased resting-state connectivity between the left lateral PFC and task-relevant regions correlated with reduced external interference (98). Therefore, in the present study, gamma activation in the lateral PFC regions, during the presentation of distracting sounds, could reflect a top-down inhibitory signal to regions involved in the processing of task-irrelevant information in young adults. Such an inhibitory signal could be supported by cortico-cortical connections of the lateral PFC to inhibitory neurons in auditory association regions (99–101).

The lateral PFC has also been proposed to control task switching, with the anterior and posterior parts of the lateral PFC playing complementary roles. The former would momentarily suspend current tasks, through inhibitory connections, and the latter would facilitate attentional switch to a novel task, through excitatory connections (review in 99). In the present study, the lateral PFC could control the shift from the TD task to the distracting sound processing, and vice versa. This is in line with a role of the lateral prefrontal cortex in the interaction between the top-down and bottom-up networks of attention (e.g. 10,102).

It is important to note that, in the present study, distracting sounds were complex sounds with acoustic spectra reaching high frequencies where older participants can display altered sensation levels. These potential differences in sensation levels between groups could result in neural activity differences. However, we believe that differences in sensation levels, related to peripheral damage, would have mostly manifested as differences in gamma activation in the ventral BU attentional network, in particular in the auditory cortices, rather than in lateral prefrontal regions. Due to the relatively low number of trials with distracting sounds and low signal-to-noise ratio of gamma activity, we were not able to factor spectral content of distracting sounds in our gamma analyses. However, the behavioral analysis on median reaction times (supplementary figure 5) according to the low or high spectral content of the distracting sounds showed neither a significant main effect of spectral content, nor a significant interaction between age and spectral content. While the absence of an effect on behavioral data does not preclude an effect on the neural data, we believe that potential differences in sensation levels is quite unlikely to explain differences in gamma activation between groups.

Therefore, the diminished activation of the lateral PFC in the elderly could result in a reduced top-down inhibitory signal, leading to both an impaired suppression of distractor processing and a delayed switching from the distracting sound back to the ongoing task, in line with an enhanced distractibility at the behavioral level.

### 4.4 Conclusion

To our knowledge, the current study is the first to utilize oscillatory modulations in the alpha and gamma bands to outline how ageing affects TD and BU attentional mechanisms in the same experiment. Behaviorally, distractibility to salient unexpected sounds is exacerbated with ageing. Modulations in alpha oscillations reveal that while facilitatory processes of TD attention seem intact, suppressive processes are reduced with ageing, showing a less efficient TD filtering of task-irrelevant information. This deficit might be compensated by an enhanced motor preparation. Moreover, modulations in the gamma activity reveal that in comparison to younger adults, elderly participants similarly activate the ventral BU attentional network but display a weaker activation, in response to distracting sounds, of frontal regions involved in inhibitory control and task-switching.

Therefore, the exacerbated distractibility exhibited by elderly participants at the behavioral level would rather be related to a reduced activation of TD inhibitory and switching processes than to an enhanced activation of the ventral BU attentional network, leading to an attentional imbalance towards an enhanced impact of BU attention at the behavioral level. This potentially frontally-driven deterioration of TD attentional control in ageing lies on the cross-roads of the two leading hypotheses accounting for the increased distractibility often observed with ageing: the inhibitory deficit (39) and the frontal ageing (40) hypotheses. In line with the proposition made by Gazzaley and D’Esposito (103), our findings reconcile both hypotheses: the decrease in the functioning of the frontal control network might be the origin of the ageing-related deficit in inhibitory mechanisms.

Finally, it is important to note that our study is limited to the attentional processes that are deployed during our paradigm i.e. TD anticipatory attention and BU attentional capture. More studies with different paradigms would be essential to test the generalizability of our results.

## Acknowledgements

We would like to thank Sebastien DALIGAULT and Claude DELPUECH for technical assistance with the acquisition of electrophysiological data. Please note that the data presented in this study have first appeared in an author’s thesis (104).

## 5 Author Contributions

HE and AB conceived and designed the study. HE and RM collected the data. HE and LF performed data (pre-)processing. HE, OB and AB wrote the paper.

## 6 Conflict of Interest

The authors declare that the research was conducted in the absence of any commercial or financial relationships that could be construed as a potential conflict of interest.

## 7 Funding

This work was supported by a grant awarded to A. Bidet-Caulet (ANR-14-CE30-0001-01) from the French National Research Agency (ANR), by a grant awarded to H. ElShafei by the Rhone-Alpes Region and by the LABEX CORTEX (ANR-11-LABX-0042). This work was performed within the framework of the LABEX CELYA (ANR-10-LABX-0060) of Université de Lyon, within the program ‘‘Investissements d’Avenir’’ (ANR-11-IDEX-0007) operated by the French ANR.

## 8 Data Availability

Data available on request. Requests should be sent to Patricia BARBIER, Laurène ALARD or Aurelie BIDET-CAULET Legal and ethical restrictions are due to the presence of potentially sensitive information e.g. MRI head shape..

## Footnotes

1 The cue by group interaction could not be directly tested because of the limitations of cluster-based permutation tests.

## Notes

### Competing Interest Statement

The authors have declared no competing interest.

